# Factors Associated with Scientific Misconduct and Questionable Research Practices in Health Professions Education

**DOI:** 10.1101/332254

**Authors:** Lauren A. Maggio, Ting Dong, Erik W. Driessen, Anthony R. Artino

## Abstract

**Introduction:** Engaging in scientific misconduct and questionable research practices (QRPs) is a noted problem across fields, including health professions education (HPE). To mitigate these practices, other disciplines have enacted strategies based on researcher characteristics and practice factors. Thus, to inform HPE, this article seeks to determine which researcher characteristics and practice factors, if any, might explain the frequency of irresponsible research practices.

**Method:** In 2017, a cross-sectional survey of HPE researchers was conducted. The survey included 66 items derived from two published QRP surveys and a publication pressure scale adapted from the literature. The study outcome was the self-reported misconduct frequency score, which is a weighted mean score for each respondent on all misconduct and QRP items. Statistical analysis included descriptive statistics, correlation analysis, and multiple linear regression analysis.

**Results and Discussion:** In total, 590 researchers took the survey. Results from the regression analysis indicated that researcher age had a negative association with the misconduct frequency score (*b* = −.01, *t* = −2.91, *p*<.05) suggesting that older researchers tended to have lower misconduct frequency scores. Publication pressure (*b* = .20, *t* = 7.82, *p*<.001) and number of publications (*b* = .001, *t* = 3.27, *p*<.01) had positive associations with the misconduct frequency score. The greater the publication pressure or the more publications a researcher reported, the higher the misconduct frequency score. Overall, the explanatory variables accounted for 21% of the variance in the misconduct frequency score, and publication pressure was the strongest predictor. These findings provide an evidence base from which HPE might tailor strategies to address scientific misconduct and QRPs.

## Introduction

In health professions education (HPE), a recent international survey of researchers reported that 90% of almost 600 respondents admitted to engaging in scientific misconduct and questionable research practices (QRPs). These practices ranged from serious infractions, like data fabrication and plagiarism, to less severe violations, like excessive self-citations, inappropriate data storage, and so-called “salami slicing” [1]. Similarly, a study of authorship in HPE unearthed a number of questionable practices, such as granting honorary authorship and denying authorship to individuals who deserved it [2]. These recent studies highlight a potential problem in HPE, a problem that can harm the research enterprise by wasting limited resources, damaging the scientific record, disadvantaging unwitting researchers, and setting a poor example for trainees [3].

To inform approaches that might mitigate research misconduct and QRPs, researchers in biomedicine [4, 5], psychology [6], and business [7] have attempted to identify relationships between personal characteristics, practice factors, and researchers’ engagement in irresponsible research behaviors. To date, these studies have considered variables such as gender, career stage, geographical location, primary research methodology used, and pressure to publish [6, 8]. For example, based on a self-report survey of biomedical researchers, Tidjink and colleagues found that publication pressure and early career stage were associated with an increased likelihood of engaging in QRPs [5]. In another study that examined authors of retracted biomedical journal articles, investigators found that junior researchers were more likely to have had a paper retracted; however, this study did not identify publication pressure as a predictor of paper retraction [8].

Although we now have a sense of the frequency of scientific misconduct and QRPs in HPE[1], we do not yet know which personal characteristics or practice factors may predict a researcher’s engagement in these irresponsible behaviors. This dearth of knowledge hampers our efforts to address misconduct and QRPs in HPE. Thus, the present study aims to address this gap by examining the associations between HPE researcher characteristics (e.g., gender, years of practice, academic background, geographic location, and primary methodology used in research); a potentially important practice factor (publication pressure); and the frequency of self-reported scientific misconduct and QRPs.

## Method

In 2017, we conducted an anonymous, cross-sectional survey of HPE researchers as a component of a larger program of research on responsible research practices. The survey’s purpose was twofold: (1) to measure the frequency of self-reported misconduct and QRPs in HPE and (2) to determine which researcher characteristics and practice factors, if any, might explain the frequency of these irresponsible behaviors. In 2018, we chose to address each of these purposes separately, as we felt that each warranted their own exploration and analysis. Therefore, we first published an article detailing the creation and execution of the survey, and we reported on the frequency of research misconduct and QRPs [1]. In this article, we now turn our attention to the relationships between researcher characteristics, practice factors, and irresponsible research practices, including both deliberate scientific misconduct and QRPs. Ethical approval for this study was granted by the Ethical Review Board Committee of the Netherlands Association for Medical Education (Dossier #937).

### Survey development

We developed a 66-item survey by adapting two published survey instruments[4, 9] (see Supplemental Digital Appendix 1). The survey items were slightly modified from the original instruments to improve clarity and relevance to the HPE context. Further details about this adaptation process, including the 19 expert reviews that were conducted, have been published previously [1].

The survey’s first set of items asked participants to indicate how often, if ever, they had engaged in research misconduct or others QRPs. The response options were *never, once, occasionally, sometimes, frequently*, or *almost always*. Participants also had the option of selecting *not applicable to my work*. In addition to questions on irresponsible research behaviors, participants responded to a nine-item, publication pressure scale, adapted from Tidjink [10], and 13 demographic items.

### Sampling and survey distribution procedures

We created our sample using two separate approaches: the first approach was based on published journal articles and the second on social media. First, we created a “curated sample” of HPE researchers that had published in 20 HPE journals between 2015-2016 (See supplemental Box 1 for listing of all journals). To identify authors, we searched these 20 journals via Web of Science, Scielo (a Latin American database), African Journals Online, and Asia Journals Online. From these articles, we extracted all author email addresses and removed any duplicate emails resulting in a sample of 1,840 unique HPE researchers. Survey invitations were distributed via Qualtrics, an online survey tool (Qualtrics, Provo, Utah), in four waves on November 13, 2017, November 20, 2017, November 27, 2017, and December 11, 2017. Of the 1,840 email invitations sent, 199 bounced back as undeliverable, reducing the number of potential respondents to 1,641.

Next, on December 11, 2017, we created the “social media sample” by posting a link to the survey to our personal accounts on Twitter and Facebook. All survey responses obtained from the social media links were tracked separately from the curated sample. Respondents in the social media sample were given the option to select “I have already completed this survey” on the survey welcome page to help us avoid duplicate responses.

### Statistical analyses

For the curated sample, we calculated the response rate based on the definitions provided by the American Association for Public Opinion Research (AAPOR) [11]. To assess potential nonresponse bias in the curated sample, we used wave analysis to calculate a nonresponse bias statistic (for additional details on this study’s application of wave analysis see Artino, 2018 [1]).

#### Measures

The outcome measure was the misconduct frequency score, which is a weighted mean score for each respondent on all 43 misconduct and QRP items. Because the misconduct and QRP items varied greatly on the dimension of “severity” (i.e., the degree to which the behavior might distort or damage science), we created a weighted frequency score using weightings from Bouter et al.’s (2016) “impact on validity” rankings [9]. For example, the item “fabricated data” had the highest severity weighting of 4.63, whereas the item “added one or more authors to a paper who did not qualify for authorship (so-called “honorary authorship”)” had the lowest severity weighting of 2.07.

The explanatory measures were the other survey variables collected, including variables measured on a nominal scale (gender, geographical region of work, academic rank, type of research, and work role) and variables measured on a ratio scale (age, publication pressure composite score, number of publications, percentage of work time doing health professions or medical education, and years involved in health professions or medical education). The publication pressure composite score was calculated as an unweighted mean score for the nine items that comprised the publication pressure scale. Selection of these explanatory variables was based on the ethical research literature from other fields [5, 6, 8].

The statistical analysis consisted of descriptive statistics of the measured variables, internal consistency reliability analysis of the nine-item publication pressure scale, correlation analysis, and multiple linear regression analysis. For the descriptive statistics, we report mean, standard deviation, and range, if a measure is on ratio scale; frequency count and valid percentage if a measure is on nominal scale. We also conducted two-way correlation analysis for the variables on ratio scale. For multiple linear regression modeling, we performed residual analysis followed by variable transformation to improve model fit. In addition, we performed dummy coding for the explanatory variables that were measured on a nominal scale. Finally, we fit a holistic linear regression model; that is, we entered all of the explanatory variables in the model at the same time. The statistical analyses were conducted in IBM SPSS 24.0 (IBM Corporation, New York, NY).

## Results

Four hundred and sixty-three respondents from the curated sample completed at least a portion of the survey, resulting in a response rate of 28.2% (potential respondents n=1,641). Based on the wave analysis, we identified a nonresponse bias statistic of 0.36. Since we utilized a six-point, frequency-response scale, this bias statistic represents a 6% difference, which, for practical interpretation of the results, is likely to have a limited effect [12].

Our social media recruitment netted an additional 127 responses. Using a multivariate analysis of variance, we compared the curated sample to those in the social media sample and found statistically significant differences *F*(5, 524) = 6.67, P < .001. Post-hoc analyses indicated that those in the curated sample were slightly older (M=47.4 years) and more experienced as HPE researchers (M=11.0 years) than those in the social media sample (M=40.7 years; M=7.5 years). However, the groups were similar in the mean number of articles published and percentage of work dedicated to HPE research. Thus, because we aimed to understand the frequency of misconduct and QRPs among a diverse, international sample, we combined the responses from the two groups and analyzed all participant responses together.

Descriptive statistics for the variables measured on a ratio scale are displayed in Table 1 and those on nominal scale reported in Table 2. Results from the reliability analysis of the nine publication pressure items indicated that the item scores had a reasonably high internal consistency reliability coefficient (Cronbach’s alpha = .83)[13].

**Table 1.** Descriptive statistics of ratio scale variables

| Variable | Mean | S.D. | Range |
| --- | --- | --- | --- |
| Weighted misconduct frequency score | .84 | .83 | 0~4.76 |
| Age | 46.03 | 11.64 | 23 ~ 87 |
| Publication pressure composite score | 2.97 | .79 | 1 ~ 5 |
| Number of publications | 40.08 | 54.98 | 0 ~ 600 |
| Percentage of work time doing health professions or medical education | 27.32 | 23.69 | 0 ~ 100 |
| Years involved in health professions or medical education | 14.91 | 9.67 | 0 ~ 54 |

**Table 2.**
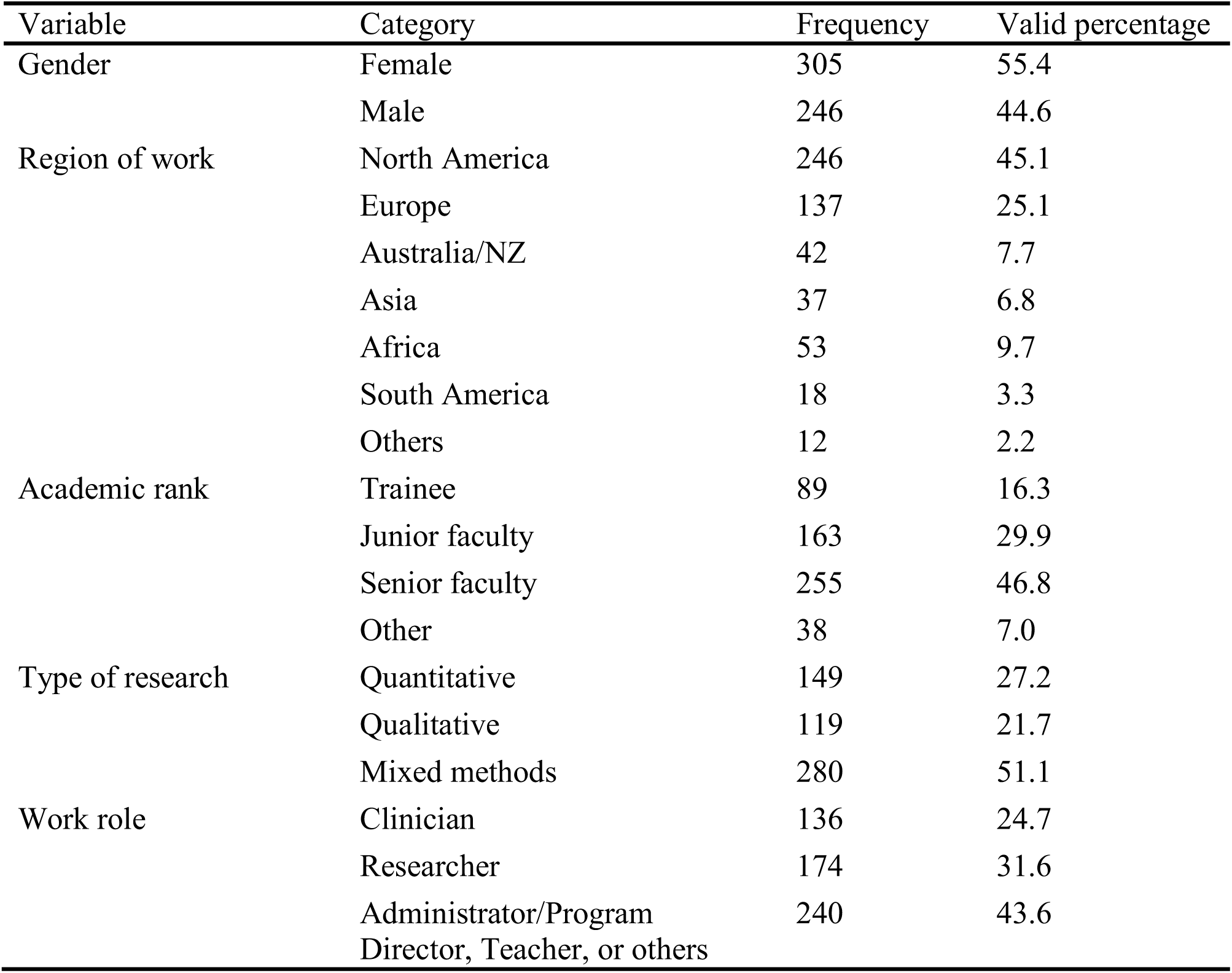
Descriptive statistics of the nominal scale variables

Results of two-way Pearson correlation analysis for the variables on a ratio scale are displayed in Table 3.

**Table 3.** Pearson correlation coefficients of the ratio scale variables

|  | Weighted misconduct frequency score | Age | Publication pressure composite score<sup>†</sup> | Number of publications | Percentage of work time doing health professions or medical education | Years involved in health professions or medical education |
| --- | --- | --- | --- | --- | --- | --- |
| Weighted misconduct frequency score |  | -.19** | .35** | .05 | .10* | -.12** |
| Age |  |  | -.34** | .48** | -.09* | .81** |
| Publication pressure composite score<sup>†</sup> |  |  |  | -.24** | .14** | -.30** |
| Number of publications |  |  |  |  | .10* | .50** |
| Percentage of work time doing health professions or medical education |  |  |  |  |  | -.05 |
| Years involved in health professions or medical education |  |  |  |  |  |  |

<sup>†</sup>The publication pressure composite score was calculated as an unweighted, mean score for the nine items that comprised the publication pressure scale (Cronbach's alpha = .83).
\* $P < .05$ ; \*\* $P < .01$

The results of the stepwise multiple linear regression are shown in Table 4. When the weighted misconduct frequency score was used as the dependent variable, the normal probability plot of standardized residual indicated unsatisfactory model fit. Thus, we transformed the variable by taking its square root. The normal probability plot of standardized residual after transformation indicated much improved model fit.

**Table 4.** Multiple linear regression modeling. (Dependent variable is the square root of weighted misconduct frequency score)

| Explanatory variable | Unstandardized regression coefficient | t-test statistic | p-value |
| --- | --- | --- | --- |
| Age | **-.01** | **-2.91** | **.004** |
| Gender | -.04 | -1.12 | .27 |
| Publication pressure and proxy of seniority |  |  |  |
| Publication pressure composite score | **.20** | **7.82** | **<.001** |
| Number of publications | **.001** | **3.27** | **.001** |
| Years involved in health professions or medical education | .003 | .84 | .404 |
| Region of work (North America as reference group*) |  |  |  |
| Europe | .03 | .54 | .591 |
| Australia/NZ | -.12 | -1.60 | .109 |
| Asia | **.21** | **2.84** | **.005** |
| Africa | .04 | .57 | .571 |
| South America | .10 | .93 | .355 |
| Academic Rank (junior faculty as reference group) |  |  |  |
| Trainee | .04 | .65 | .519 |
| Senior faculty | -.004 | -.09 | .929 |
| Type of research (quantitative methods as reference group*) |  |  |  |
| Qualitative | .06 | 1.15 | .249 |
| Mixed-methods | -.02 | -.56 | .577 |
| Work role (clinician as reference group) and percentage of time doing research |  |  |  |
| Researcher | **.12** | **2.15** | **.032** |
| Other work role | .03 | .60 | .550 |
| Percentage of work time doing health professions or medical education | -.001 | -.72 | .474 |
| Total Model $R^2$ | | .21 | |

\*These groups were selected as the reference groups because they represented the largest respondent groups in their respective categories.

Researcher age had a significant negative association with the misconduct frequency score (*b*= −.01, *t*=-2.91, *p*<.05). This suggests that older researchers tended to have lower misconduct frequency scores. Publication pressure (*b*=.20, *t*=7.82, *p*<.001) and number of publications (*b*=.001, *t*=3.27, *p*<.01) both had significant positive associations with misconduct frequency score. The greater the reported pressure or the more publications listed, the higher the misconduct frequency score. Researchers from Asia showed a relatively higher misconduct frequency score compared to those from North America (*b*=.21, *t*=2.84, *p*<.01). The work role of “researcher” was associated with a higher misconduct frequency score compared to the work role of “clinician” (*b*=.12, *t*=2.15, *p*<.05). Overall, these results indicate that publication pressure was by far the strongest predictor of the misconduct frequency score. Taken together, all of the explanatory variables explained 21% of the variance in misconduct frequency. It should be noted that all the unstandardized regression coefficients reported in this paper were obtained by using the square root of weighted misconduct frequency score as the dependent variable in the regression model.

## Discussion

This analysis of HPE researchers’ self-reported engagement in scientific misconduct and QRPs suggests that the variables of age, publication pressure, number of publications, geographical location, and work role explain considerable variance in the frequency of researchers’ self-reported irresponsible research behaviors. Taken together, these findings provide the field with an evidence base from which to tailor strategies for addressing scientific misconduct and QRPs in HPE.

Our findings indicate that older HPE researchers report misconduct and QRPs less frequently than young researchers. These results provide some evidence for the idea that junior scientists may be particularly vulnerable to the lure of misconduct and QRPs in HPE or, alternatively, that junior researchers may simply be unfamiliar with responsible research practices [8]. In the biomedical and social sciences, this finding has led funders, such as the National Institutes of Health, to mandate that all grantees, including junior and senior researchers, complete responsible conduct of research training [14-16]. While such an approach is a positive step, it is worth nothing that most HPE research is unfunded, making it possible that our junior researchers may not be offered or exposed to such training. Furthermore, researchers have wrestled with the scope of this training, raising concerns that it often falls short of addressing the increasingly diverse nature of research [14, 17, 18]. As noted by Keune et. al., when studying common HPE topics, like examining residents and trainees, this training’s lack of diversity can mean many researchers are operating without guidance on how to navigate common ethical challenges within HPE [19].

Researchers have long described the pressure to publish as a contributor to the “dark side of science” [20]. In biomedicine, researchers experiencing publication pressure are more likely to admit engaging in misconduct and QRPs [4, 8, 21, 22]. The findings reported here align with and extend this previous work. We found that both perceived publication pressure and the number of publications were associated with misconduct and questionable research conduct. To mitigate the pressure to publish, researchers have suggested modifying promotion and tenure structures, promoting and rewarding research transparency, and training senior researchers to act as responsible role models by demonstrating positive (and ethical) research practices [8, 23]. Integrating such approaches in HPE may be beneficial; however, in many cases, HPE researchers are not directly assigned to a department of HPE, but rather are often members of clinical departments (e.g., Departments of Medicine, etc.). Therefore, for role modeling to have a positive impact of HPE researchers, role such practices would likely need to be integrated at the institutional level.

To our knowledge, this is the first study to directly document that HPE researchers feel pressure to publish. Not only has the pressure to publish been connected with greater misconduct and QRPs, but it has been tied to researchers’ reluctance to share their work openly and to partner with other researchers. It has also been linked to higher levels of researcher burnout [24, 25]. Overall, these findings suggest that the topic of publication pressure may be ripe for further exploration in HPE.

Our findings suggest that HPE researchers in Asia reported higher frequencies of irresponsible behaviors than researchers in North America. In the wake of concerns that research misconduct in Asia is on the rise, multiple research teams have undertaken Asia-specific programs of research [26] to further examine the problem. For example, a team of Chinese scientists recently conducted a survey and found that 40% of research from China may be tainted by researcher misconduct [27]. In that particular study, the respondents attributed misconduct to lack of training and institutional oversight, and to extremely high pressures to publish. In studying geographical influence more broadly, Fanelli suggests that geographical variation in research misconduct may be related to the presence or absence of policies and procedures [8].

Respondents identifying primarily as researchers reported higher frequencies of QRPs and misconduct than those identifying as clinicians. Although speculative, this finding could be related to the idea that the stakes tend to be much higher for those participants for whom research output and published papers are the primary measures of success. On the other hand, a clinician who conducts education research “on the side,” may be less incentivized to cut corners. Regardless of the explanation, we would advocate for future research to more closely explore this relationship. Related areas of exploration could include the impact of widespread and long-standing ethics training in clinical education [28, 29] in contrast to approaches taken (or not taken) by graduate programs in HPE. Future studies might also consider differences in the incentive structures and other systematic pressures for researchers as compared to clinicians.

### Limitations

Our study has several limitations. The curated sample had a modest response rate of 28.2% and therefore nonresponse bias is a real possibility. Although the wave analysis indicated that nonresponse bias likely had limited effects on our results, it is possible that certain groups, (e.g., senior researchers) are over-represented in our sample, thereby biasing our findings.

Responsible research behavior is complex and context specific. We recognize that measuring such complex phenomena with self-report survey items has limitations. Moreover, self-report surveys can be sensitive to several types of response bias, including limitations of autobiographical memory and differences in interpretations of the meaning of questionable research behaviors. Notwithstanding these limitations, our findings align with the broader literature on ethical research practices. For example, Fanelli et. al. conducted a meta-analysis of 21 survey studies and concluded that 2% of respondents admitted to fabrication, a finding that exactly matches our results for this particular type of misconduct [30].

Lastly, we recognize that an individual’s decision to engage in research misconduct and/or QRPs is complex. Therefore, while we have identified a set of variables that seem to explain a fairly large proportion of variance in these behaviors, an even larger proportion (almost 80%) is still unexplained. What is more, the correlational nature of this study precludes us from making strong statements regarding causality; it also prevents us from making definitive suggestions for how we might mitigate such practices.

## Conclusions

The results of this study suggest that a researcher’s age, number of publications, geographical location, and work role are associated with the frequency of self-reported misconduct and QRPs in HPE. We also found a positive association between perceived publication pressure and irresponsible research behaviors. Both scientific misconduct and QRPs have the potential to seriously damage the quality of our scientific work, as well as the public’s belief in its credibility. Ultimately, we hope these results can help academic leaders, journal editors, and others involved in educational research develop targeted measures to minimize scientific misconduct and QRPs in HPE.

Declaration of interest

Erik Driessen is the current editor-in-chief of *Perspectives on Medical Education*. He was not involved in the review of or the decision to publish this article.

The views expressed in this article are those of the authors and do not necessarily reflect the official policy or position of the Uniformed Services University of the Health Sciences, the U.S. Navy, the Department of Defense, or the U.S. Government.

## Acknowledgements

The authors acknowledge and thank all of the HPE researchers that completed the survey.

